# Expansion of the genomic and functional diversity of global ocean giant viruses

**DOI:** 10.1101/2024.07.26.605358

**Authors:** Benjamin Minch, Mohammad Moniruzzaman

## Abstract

Giant viruses (GVs) play crucial roles in the global ocean microbial food web and biogeochemistry by infecting protists. Traditionally, insights into GV ecology and functions have been limited to culture-based studies. However, recent metagenomic advances have uncovered over 1,800 new GV genomes from the world’s oceans. While this rapid increase in genomic information marks an impressive advance for the field, it is nowhere close to the extensive genomic information available for other marine entities - e.g., prokaryotes and their ‘virome’. Addressing this gap, we present 230 novel, high-quality GV genomes and 398 partial genomes from nine global ocean datasets using an open-source bioinformatic workflow we developed. Notably, we identified numerous GV genomes from the Baltic Sea, offering insights into their phylogenomics, metabolic potential, and environmental drivers in one of the largest brackish water ecosystems. We discovered new GV functions, including photosynthetic proteins, and identified a significant functional divide between the Imitervirales and Algavirales orders. Additionally, we evaluated best practices for GV genome recovery from metagenomic datasets through a case study on the Baltic Sea dataset. Our study significantly expands the marine GV genomic and functional diversity, broadening our understanding of their roles in the food web and biogeochemistry.

## Introduction

The discovery of giant viruses (GVs) belonging to the phylum *Nucleocytoviricota* represents a paradigm shift in the world of virology due to their remarkable genomic and functional complexities [1]. These viruses, with their large virion (up to 2uM) and genome sizes (up to 2.5Mbp)[2,3] have redefined our understanding of viruses. Of note is their ability to encode numerous functionalities previously unknown to viruses, which has raised significant questions regarding their ecological and evolutionary roles in the biosphere [4–6]. Currently, six identified orders exist for GVs - including Imitervirales, Algavirales, Pimascovirales, Asfuvirales, Pandoravirales, and Chitovirales [7]. Members of several of these orders have a seemingly broad host range, consisting of mainly single-celled eukaryotes [8,9] and are widespread in the global oceans and sediments [10,11].

The impact of giant viruses is no doubt present across all environments, but special interest has been taken to uncover their impact on aquatic ecosystems due to the important contribution of single-celled eukaryotes to global biogeochemical cycles and aquatic food webs [12,13]. Giant viruses infecting these cells have been shown to modulate host metabolism during infection [14–16] through the introduction of a diverse set of metabolic genes encoded in the GV genomes [4,5]. Giant viruses also have the potential to modulate how hosts acquire nutrients through the use of viral transporter proteins [17,18]. This impact on host metabolism has potential implications for marine nutrient cycling [19] and food webs [20].

Ever since the discovery of the first giant virus in the 1980s [21] many methodological and technological strides have facilitated our understanding of the diversity and functionality of these viruses. Much of the early understanding of GVs came from culture-based approaches [22] and the first GV genome wasn’t sequenced until 2004 [23]. While this culture-based approach is invaluable in the understanding of GV biology and there have been many advances in virus recovery methods [24,25], to date, there are only around 200 genomes recovered from isolates [26]. Compared to the over 14,000 bacteriophage genomes [27] and 600,000 bacterial genomes [28], this number is minuscule.

In recent years, this culture-based approach has been complemented by large-scale metagenomics to find signatures of GVs in environmental data. Early metagenomic methods relied on the recovery of GV signatures such as polymerases and major capsid proteins [10,29–34]. These early surveys demonstrated the widespread distribution of GVs and suggested there was a huge pool of undiscovered diversity of these viruses [35]. The first giant virus Metagenome-Assembled Genome (GVMAG) was recovered in 2011 from the Organic Lake, Antarctica [36]. This recovery was followed by an additional four genomes recovered from Yellowstone Lake [37] as well as a few more genomes derived from wastewater, soil, lake, and deep-sea sediment samples [11,38–40]. The power of metagenomic-based approaches became most evident with the discovery of 2,074 GVMAGs from diverse environments [41], and the unveiling of 501 GVMAGs from mostly marine environments [42]. In the years to follow, only one other major study has recovered a large diversity of additional GVMAGs from marine environments, focusing on the TARA global oceans data to build the Global Ocean Eukaryotic Viral database (GOEV) [43]. In total, Gaïa et al. discovered over 400 additional GVMAGs as well as expanding diversity to include the putative new class *Proculvirales* and the new phylum *Mirusviricota,* which reveals evolutionary connections of giant viruses to herpesviruses.

While much progress has been made in recovering GVMAGs from metagenomic datasets, we still have far to go in terms of recovering additional diversity and functional potential of giant viruses, especially their diversity in the oceans as only ∼1800 genomes exist from marine datasets [43]. Further efforts towards understanding GV diversity and functional potential in the global ocean are necessary for a comprehensive assessment of their impact on marine protist ecology and biogeochemical cycles. To this end, we developed PIGv, an open-source, easy-to-use bioinformatic pipeline which should facilitate GV genome discovery from diverse environmental metagenomic datasets. Leveraging this pipeline, we report 230 novel high-quality marine GVMAGs and 398 partial GVMAGs from 9 different datasets throughout the global oceans. Analysis of these data reveals novel functional potential encoded by giant viruses and their ecological constraints and provides insights into best practices in methodologies for GVMAG recovery. We also reveal a large number of GVMAGs from the Baltic Sea, an ecologically unique, large brackish water body for which little data exists on the phylogenetic diversity and ecology of GVs.

## Methods

### Data Acquisition

To recover giant virus genomes, we downloaded raw sequencing data (illumina) from 9 publicly available BioProjects which represent 8 different bodies of water spanning from pole to pole (Table S1). These projects were chosen based on their sampling of the cellular size fraction (>0.2uM) and the high number of hits to giant virus major capsid proteins (MCPs) that we assessed using a pre-screening tool we previously developed (https://github.com/BenMinch/PIGv/tree/main/viral_screening). We also avoided large datasets that had already been searched for GVs such as the Tara Oceans data [43]. All raw metagenomic reads from these projects were downloaded and trimmed with trimGalore using default parameters [44]. The raw reads were then assembled using Megahit [45] and reads were mapped back onto the assemblies to generate coverage using CoverM (https://github.com/wwood/CoverM) in coverage mode.

### Genome Recovery and the PIGv pipeline

We developed PIGv (Pipeline for Identification of Giant Viruses), a bioinformatic pipeline to facilitate the assembly of giant viruses from metagenomic libraries (Figure 1). PIGv combines several existing software and scripts to facilitate this process, along with newly developed screening and annotation scripts. Assembled contigs and coverage files were used as inputs for PIGv (github.com/BenMinch/PIGv). This tool first bins metagenomic contigs using metabat2 [46] and then predicts proteins for bins using prodigal-gv [47,48] which is a version of prodigal trained with giant virus models and alternative genetic codes. Bins are then screened using the NCLDV markersearch script [7] which searches for key marker genes present in giant virus genomes (DNA polymerase family B (PolB), DNA-directed RNA polymerase beta and alpha subunits (RNAPS, RNAPL), NCLDV major capsid protein (MCP), Packaging ATPase (A32), Poxvirus Late Transcription Factor (VLTF3), DNA topoisomerase II (TOPOII), DEAD/SNF2-like helicase (SFII), and Transcription initiation factor IIB (TFIIB)). Bins that had at least one marker gene were kept for further screening.

**Figure 1.**
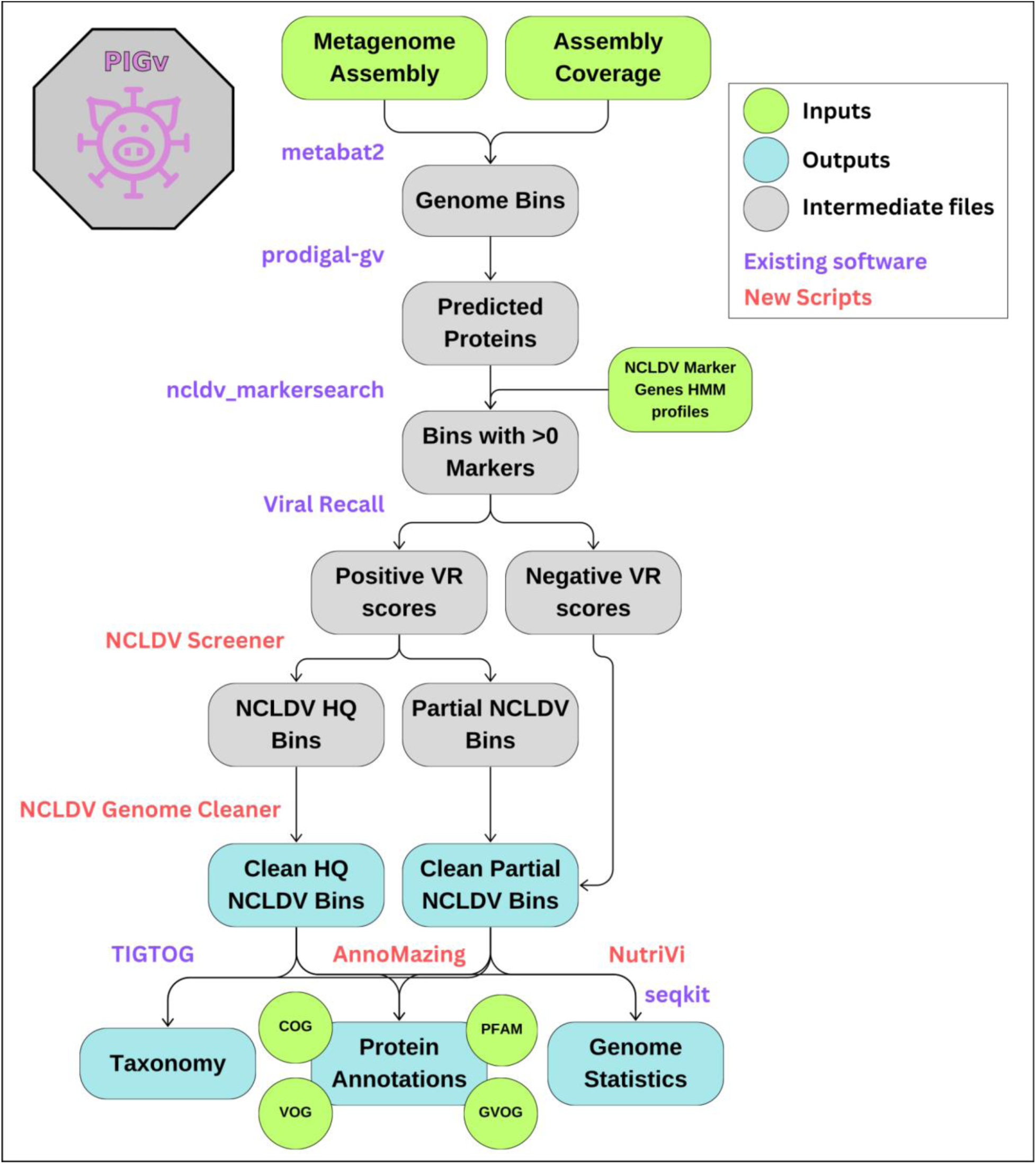
Pipeline for identification of Giant viruses (PIGv). The entire binning and screening pipeline was developed using existing (purple) and newly developed (red) scripts, taking a metagenome assembly as input. The automated pipeline produces both high-quality (HQ) and partial Giant virus bins as well as annotations, taxonomy, and statistics for those bins.

After running a search for marker genes, the bins with at least one marker gene were screened using ViralRecall [49] to confirm they were giant viruses. Bins with negative ViralRecall scores (meaning they most likely weren’t giant viruses) were discarded. After this screening, another screening was done to keep “high-quality” giant virus bins. Bins were only kept if they had at least 3 of the following marker genes (PolB, MCP, A32, VLTF3, SFII). These bins were then processed using ViralRecall in contig mode, which screens individual contigs within the bins and removes those with negative scores. Putative Mirusviricota bins were also screened using this pipeline and were kept if they contained a hit to a Mirus MCP (using a custom hmm profile) and at least one other Mirursviricota marker gene (RNA pol A, RNA pol B, DNA pol B, TFIIS).

Partial giant virus bins were also recovered by taking any bins with at least one marker gene and screening them through ViralRecall in contig mode to remove non-giant viral contigs. These “clean” partial bins were used for subsequent protein analysis.

All genomes were dereplicated at 98% ANI from the Global Ocean Eukaryotic Virus database (GOEV) with a 25% minimum coverage threshold using dRep [50]. Cleaned partial bins were dereplicated at 99% ANI with a 25% minimum coverage threshold to get rid of duplicate bins. This resulted in a total of 230 new giant virus genomes and 398 cleaned partial bins for downstream analysis.

Basic genome statistics such as length and GC percentage were calculated using seqkit [51] and nitrogen/sulfur per sidechain were calculated using a script adapted from [52](github.com/BenMinch/NutriVi). Comparisons of nitrogen and sulfur per sidechain, average length, and GC content between orders were done using a two-sample t-test. All plots were made using ggplot2 [53] in R.

### Taxonomy and Phylogeny

Taxonomy was predicted for all genomes using TIGTOG [54], a recently developed tool that classifies giant virus genomes or partial genomes based on protein family trademarks. In addition to classifying genomes, a phylogenetic tree was made using the PolB marker gene. Briefly, PolB genes were gathered from all the high-quality and partial genomes and dereplicated at 98% using cd-hit [55]. These proteins were aligned with a curated set of reference PolB representing all major giant virus families from the GOEV database [43]. Proteins were aligned using MAFFT (--auto) [56] and alignments were trimmed using trimAL with the ‘gt 0.1’ parameter [57]. A maximum-likelihood phylogenetic tree was made using IQ-TREE [58] with the ‘LG+F+R10’ model with 1000 bootstraps. The tree was visualized in iTOL [59].

### Protein Annotation

All proteins from both quality and partial bins were predicted using prodigal-gv. These predicted proteins were then annotated with the PFAM [60], GVOG [7], VOG [61], and COG [62] databases using Annomazing (github.com/BenMinch/AnnoMazing). This tool performs annotations based on hmm searches against requested databases and compiles them all together. For all searches, an e-value cutoff of 1e-5 was used. Genomes in the entire GOEV database were also annotated similarly.

Proteins that are unique to the genomes generated in this study were identified by first clustering our new genome-encoded proteins using mmseqs2 with 50% identity (coverage mode 5, -c 0.4) [63]. After clustering, representative sequences from each cluster were searched against a database of all GOEV (Global Ocean Eukaryotic Virus) and giant virus database (GVDB) [7] proteins using BLAST. Proteins without a significant hit (e-value greater than 1e-3) were chosen for further analysis. These representative proteins were then screened for unique functional annotations by looking for unique PFAM accession numbers not present in GOEV annotated genomes. This yielded a total of 569 novel protein clusters in our genomes.

Metabolic genes for high-quality genomes were annotated using NuMP (Nucleocytoviricota metabolic profiler) (github.com/BenMinch/NuMP), which looks for specific metabolic genes that have been found previously in giant viruses. These were displayed as a heatmap using pheatmap (github.com/raivokolde/pheatmap) based on presence/absence. Functional differences between Imitervirales and Algavirales genomes were displayed using the ‘anvi-display-functions’ program in Anvi’o (v8) with COG annotations [62,64].

### Assessment of GV abundance in the Baltic Sea metagenomes

Abundance profiles of high-quality GVMAGs recovered from the Baltic Sea (n=108) were acquired through mapping trimmed reads from the Baltic Sea metagenomes to the genomes at 95% ANI using CoverM with minimap2 [65]. Read counts were normalized by library and genome size and used to access GV abundance in different size fractions as well as correlations with various environmental variables.

To assess correlations between giant virus genome abundance and environmental variables, a Mantel test was performed using Spearman correlation and 9999 bootstraps in the vegan R package (github.com/vegandevs/vegan). A canonical correlation analysis (CCA) was also done on individual genomes using the same environmental variables.

## Results

### Biogeography of 230 novel giant virus genomes

After applying our giant virus recovery pipeline, 230 high-quality non-redundant giant virus metagenome-assembled genomes (GVMAGs) were recovered from 9 different datasets (See methods)(Table S1). These datasets encompassed 4 of the 5 major oceans with a special focus on colder bodies of water (Figure 2a). The most high-quality GVMAGs were recovered from the Baltic Sea (n=108), followed by the Antarctic (n=65).

**Figure 2.**
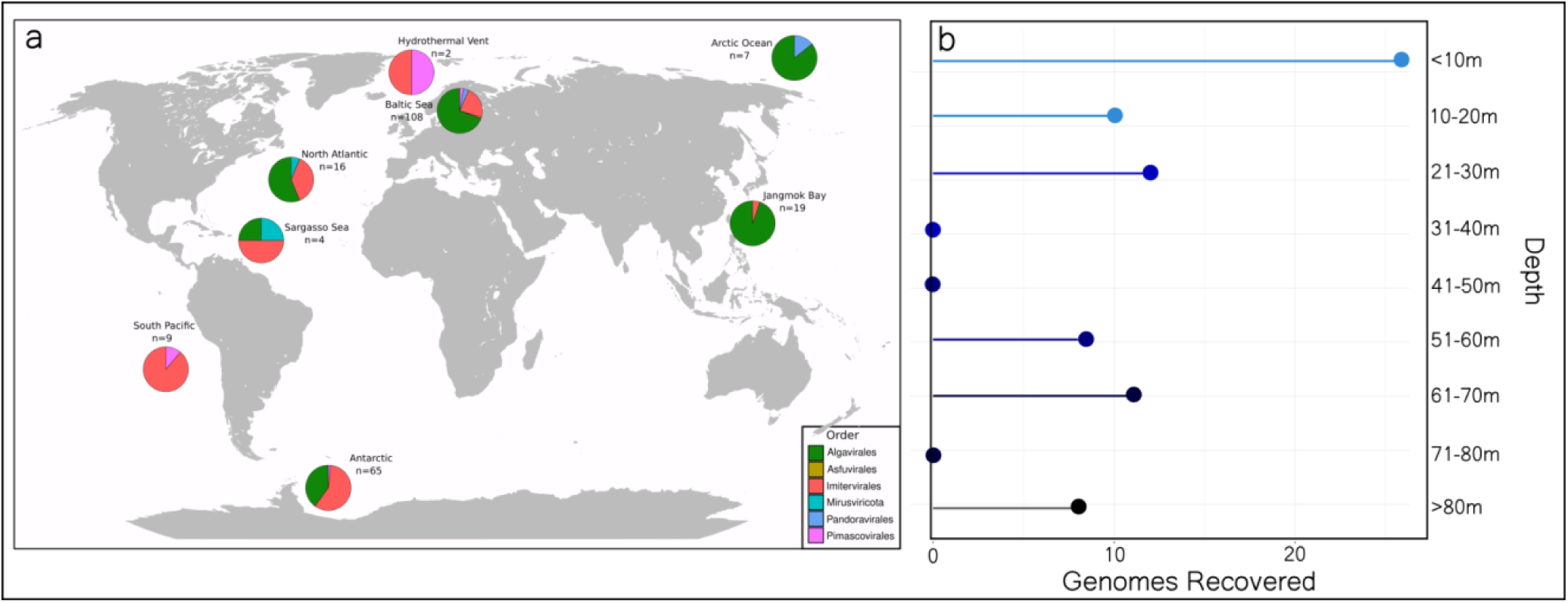
Global distribution of recovered giant viruses. (a) Giant virus metagenome-assembled genomes (GVMAGs) of every major giant virus order were acquired from datasets spanning pole to pole. The pie charts represent the proportion of genomes recovered from each order from each locale. (b) For sequencing projects with available depth data, this information was used to show recovery success at various depths.

We recovered giant viruses from all major orders, with the majority of them from either Algavirales (n=135) or Imitervirales (n=81) orders. Two novel Mirusviricota genomes were also recovered from the Sargasso Sea and North Atlantic datasets. Although not all datasets had accompanying depth data, the ones that did show that most of these GVMAGs originated from the sunlit surface oceans except for 2 coming from a hydrothermal vent environment (Figure 2b).

In addition to the 230 high-quality GVMAGs, 398 partial GVMAGs were also recovered from the datasets (Figure S1). The majority of these came from the Baltic Sea (n=203) and were classified as either Algavirales (n=127) or Imitervirales (n=102).

### Genome Statistics

Recovered GVMAGs ranged in length from 50 kbp to 1.3 Mbp, averaging 211 kbp (Figure 3). Broken down by order, Algavirales GVMAGs were significantly larger than other orders with an average length of 362 kbp. This larger average size is likely due to the three largest GVMAGs belonging to the Algavirales order.

**Figure 3.**
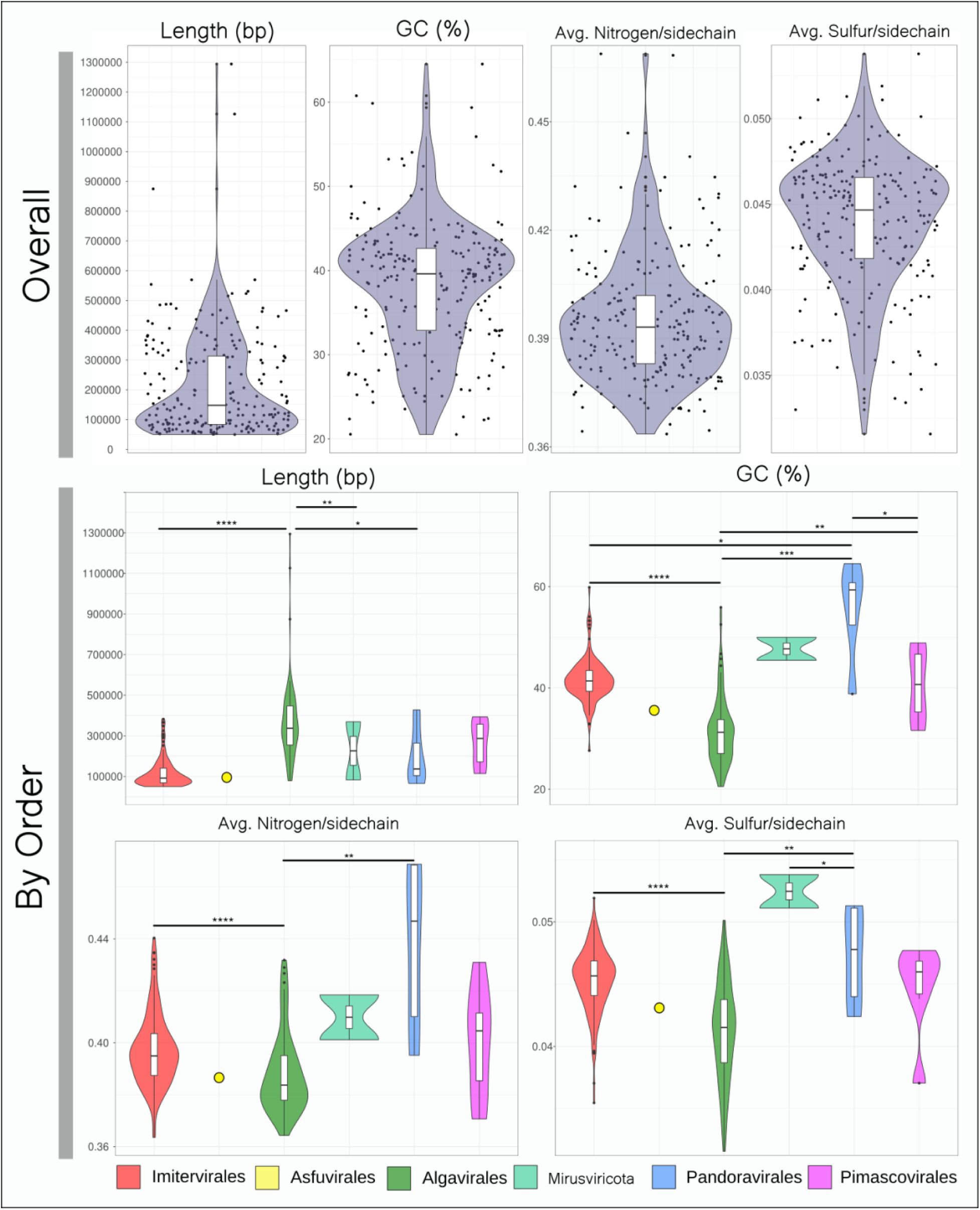
Genomic information of recovered GVMAGs. Violin plots representing length, GC percentage, average nitrogen per sidechain, and average sulfur per sidechain were constructed to show differences between GV phylogenetic orders. For each order with more than 2 recovered genomes, a t-test was performed to see statistically significant differences in these genomic characteristics. (*= p<.05, **= p<.01, ***= p<.001, ****= p<.0001).

The average GC percentage for all recovered GVMAGs was 38.37%. This average did have significant differences between orders as Pandoravirales had the highest average GC percentage at 55%. Algavirales had on average the lowest GC percentage with 31.5%.

To obtain further insights into the protein makeup of the GVMAGs, we calculated the average nitrogen per sidechain and average sulfur per sidechain for all proteins in the GVMAGs. On average these numbers varied only slightly between genomes with an average of 0.395 nitrogen molecules and 0.044 sulfur molecules per sidechain. Significant differences did exist between orders as Pandoravirales had the highest average nitrogen per sidechain (0.44) and Mirusviricota genomes had the highest average sulfur per sidechain (0.052).

### Phylogenetic analysis

A phylogenetic reconstruction of the newly discovered high-quality and partial GVMAGs based on the DNA Polymerase B marker gene (PolB) showed that many of these GVMAGs clustered within known giant virus families (Figure 4). Phylogenetic analysis did not reveal biogeography to be a key driver of the evolutionary history of these genomes, as genomes from diverse orders were recovered from different oceanic regions without any location-specific clustering on the tree.

**Figure 4.**
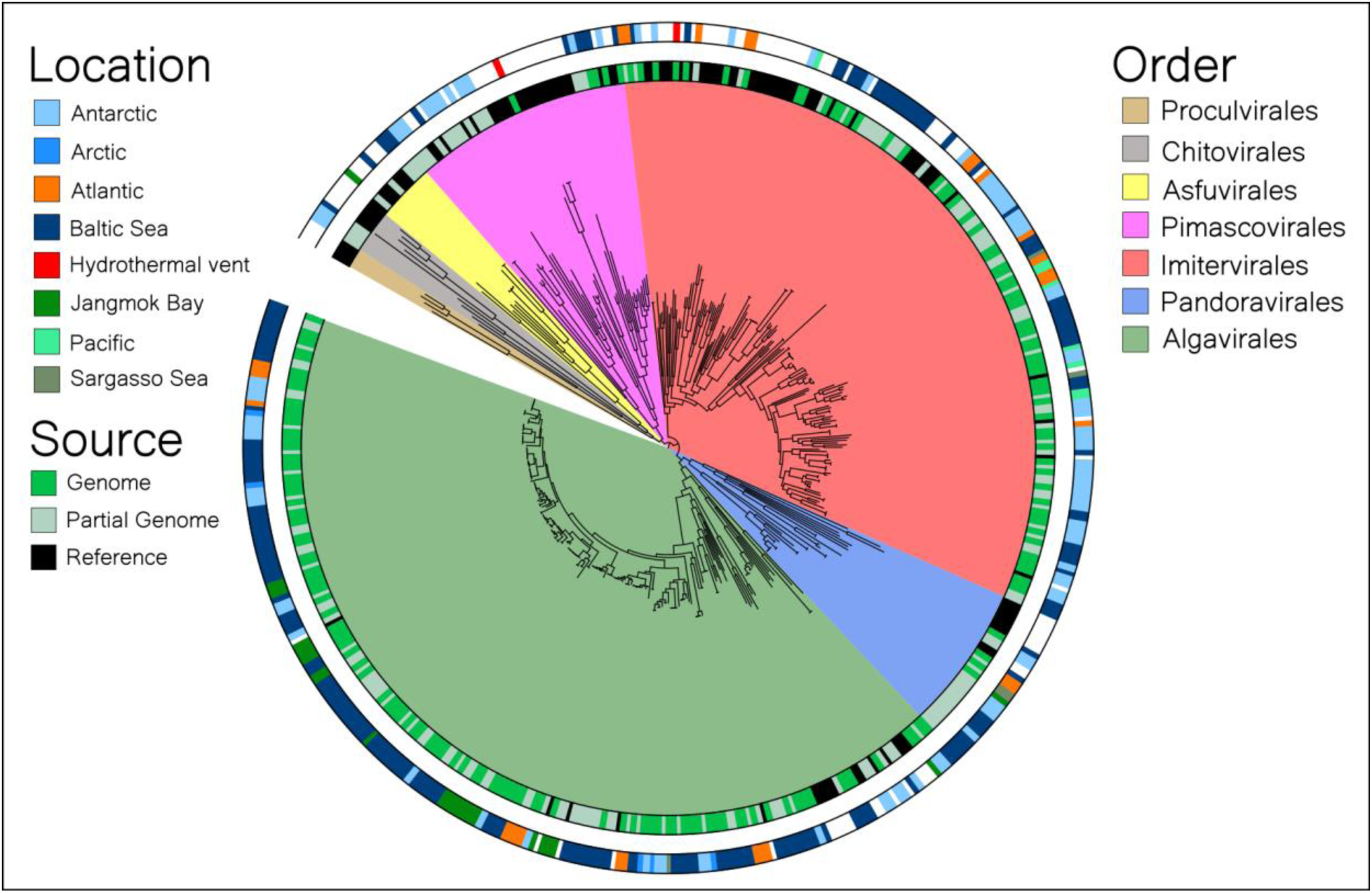
Phylogenetic placement of recovered GVMAGs. A phylogenetic tree was constructed using the PolB marker gene from our recovered GVMAGs as well as PolB genes from a curated set of reference genomes representing all major GV orders and families. The type of sequence as well as the location of recovery are represented as colorstrips on the tree. The tree was constructed using IQ-TREE with the ‘LG+F+R10’ model and 1000 bootstraps. It was visualized using iTOL.

### Phylogeny-informed metabolic potential of Giant viruses

#### Central Carbon Metabolism

Genes involved in central carbon metabolism (Glycolysis, TCA cycle, and the pentose phosphate pathway) were commonly found in Imitervirales genomes with some genomes even having multiple genes involved in the pathway. Despite larger genome sizes of recovered Algavirales GVMAGs, central carbon metabolism genes were nearly absent from these genomes with glycolysis genes being present in only 3 genomes, phosphate pentose genes being present in 7 genomes, and TCA cycle genes being completely absent.

#### Light harvesting and Beta Oxidation

The light-harvesting Chl a/b-binding (LHCB) protein was found across orders at similar rates of occurrence except for it being absent in recovered Pimascovirales genomes. On the other hand, rhodopsins were found exclusively in recovered Imitervirales genomes, showing a stark phylogenetic separation. These proteins were quite common within this order, being found in 53% of recovered genomes.

Genes involved in beta-oxidation also showed order-level patterns, only found in recovered Pimascovirales (20%) and Imitervirales genomes (25%).

#### Nutrient Transporters and Metabolism

Certain genes involved in nutrient metabolism and transport also showed order-specific bias in genomic-distribution. Specifically, no Algavirales or Pandoravirales genomes contained genes for sulfite export while Imitervirales (25%) and Pimascovirales (20%) recovered genomes had these genes. Genes for nutrient metabolism such as Glutaminase, Glutamine synthetase, and Pho regulon protein were also absent from Algavirales and Pandoravirales genomes. These genes were only found in recovered Imitervirales and Pimascovirales genomes with Glutamine synthetase being found in 22% of Imitervirales genomes and Pho regulon protein being found in 40% of Pimascovirales genomes.

#### DNA processing

Genes involved in DNA processing were found almost exclusively in recovered Imitervirales genomes. A surprising 95% of these genomes were found to contain a DNA mismatch repair gene (MutS) while all other orders had much less representation of this gene (5% in Algavirales; 20% in Pimascovirales; 11% in Pandoravirales). Histone acetyltransferase genes were also found in 20% of Imitervirales and 20% of Pimascovirales genomes while no Algavirales or Pandoravirales genomes had them.

#### Broad functional divide between Imitervirales and Algavirales genomes

Beyond differences in key metabolic genes, the two major orders of giant virus recovered in this study show broad functional differences encoded in their genomes (Figure 5b). Out of the 575 COG (clusters of orthologous genes) annotations from both groups of genomes, 383 (67%) were found to be exclusive to Imitervirales, 40 (7%) to Algavirales, and 152 (26%) were shared between the two groups.

**Figure 5.**
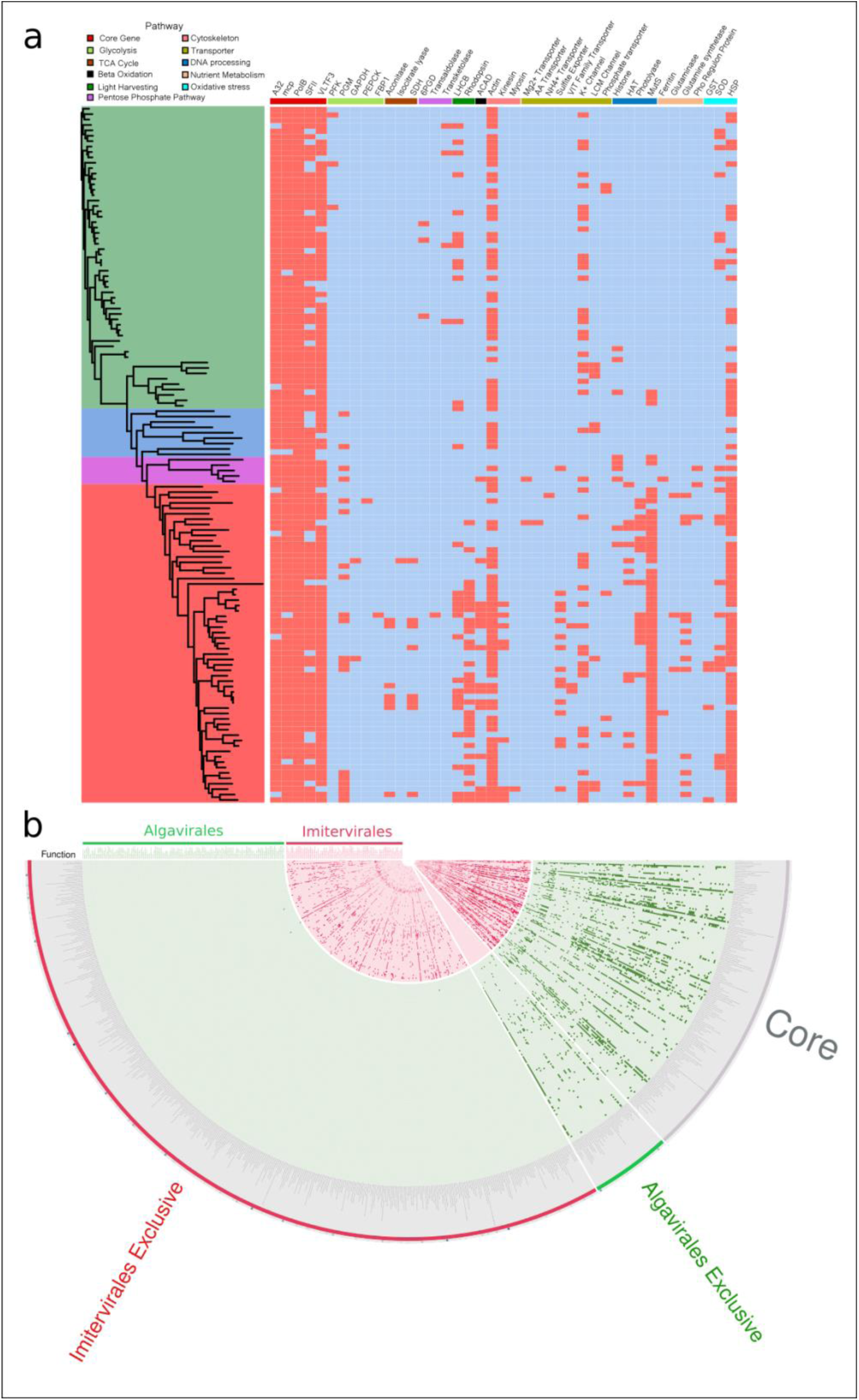
Phylogenetically informed metabolic potential of recovered GVMAGs. (a) A heatmap showing the presence(red)/absence(blue) of common giant virus metabolic genes in our recovered GVMAGs. These genes are separated into pathways based on cellular function and are organized into phylogenetic groups based on the phylogeny of their PolB (see previous figure). (b) Functional differences between Imitervirales and Algavirales genomes visualized in Anvi’o. Each ring represents a genome, and each column is an individual COG functional annotation. The plot is broken up into sections based on functions present only in Imitervirales genomes, those present only in Algavirales genomes, and those shared between the two as core genes.

### New functionality found in GVMAGs and partial MAGs

A survey of the functional landscape in our newly recovered GVMAGs and partial GVMAGs revealed 569 novel proteins not found previously in giant viruses after dereplication of protein clusters and annotations from both the Global Ocean Eukaryotic Virus (GOEV) database and the Giant Virus Database (GVDB) (Figure 6a). Most of these novel proteins have unknown GO functional annotations, with many also having a role in protein binding, membrane interactions, and ATP binding (Figure 6b). The vast majority of the new functions are encoded within genomes from the Arctic, Antarctic, and Baltic Sea, contributing to 88% of the total new proteins.

**Figure 6.**
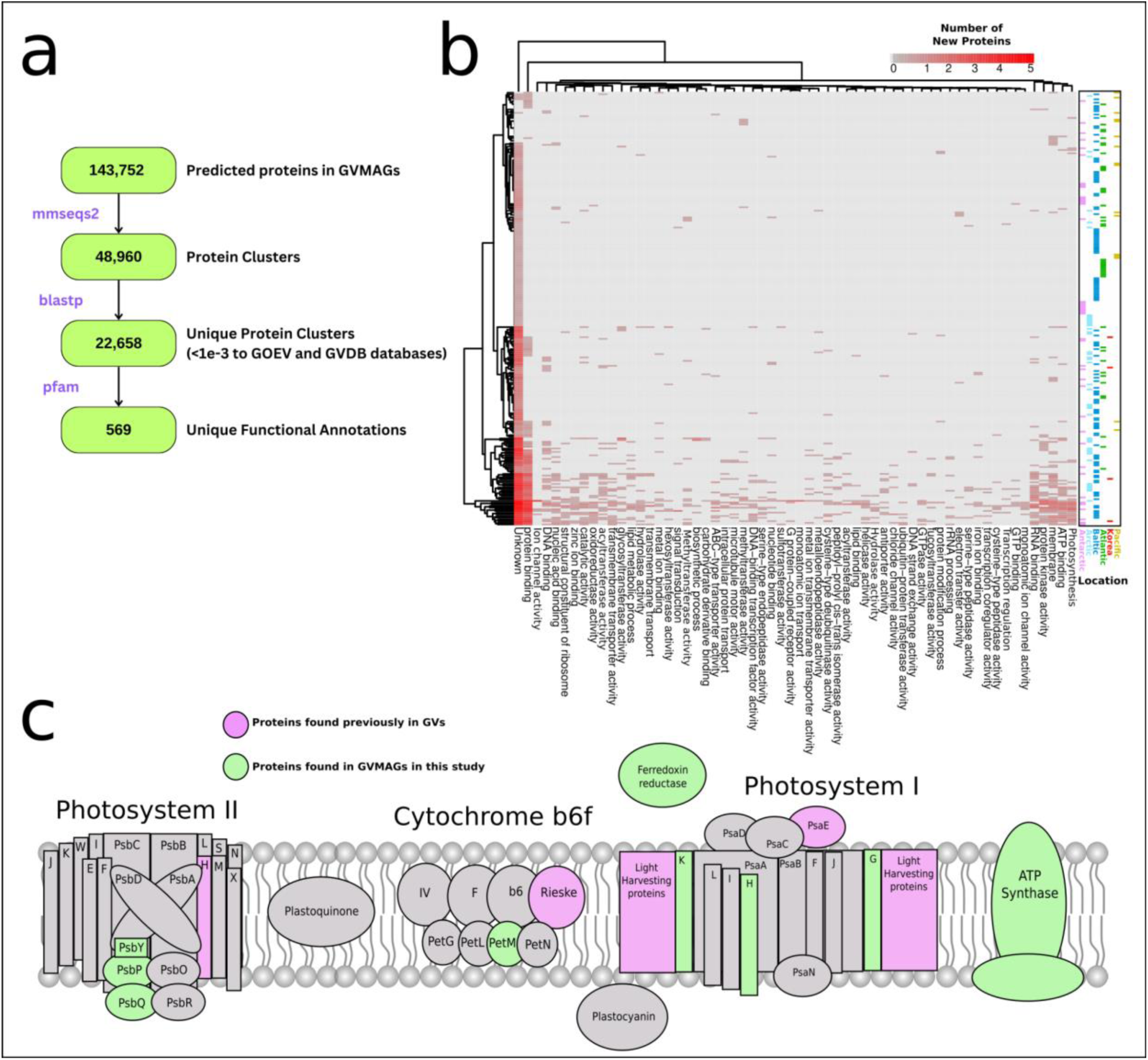
Unique functional potential in recovered GVMAGs. (a) Proteins that have never been found in giant viruses (dereplicated from the GOEV and GVDB databases) were annotated to expand the functional potential of known GVs. (b) A heatmap of unique functional potential in new GVMAGs. Each row represents a genome and columns are separated by gene ontology (GO) categories. The location of the recovered genome is also shown on the right. (c) Specific unique genes involved in the photosynthesis pathway are highlighted here and combined with proteins found in previous studies such as those by Schultz et al. [41] and Gaia et al [43].

This survey also revealed many new proteins putatively involved in photosynthesis such as those involved in photosystem I (PsaK, PsaG, PsaH, Ferrodoxin reductase), photosystem II (PsbQ,PsbY, PsbP), cytochrome b6f (PetM) and ATP synthase (Figure 6c). This greatly expands the number of known photosynthetic genes encoded in giant virus genomes as reported in other studies [41,42].

### A case study on factors affecting GV recovery and abundance

Due to the large number of Baltic Sea samples, as well as the experimental design utilizing multiple filter sizes, we were able to utilize this data to perform a case study on factors affecting genome recovery, size, and abundance. Each sample in the Baltic Sea dataset used one of 5 different filtering ranges, all within the plausible size range for the expected recovery of giant virus genomes. Between the 5 size ranges, the most average recovered genomes per sample came from the 0.1-0.8 um range, followed by the 0.8-3 um range (Figure 7). The lowest average number of genomes were recovered from 3-200um size ranges. Genomes from the >0.2 um size range had the largest average genome size as well as the largest range in genome sizes, while the 0.8-3 um range had the lowest average recovered genome size. In terms of reads mapped to recovered GV genomes, the larger size ranges (0.8-3.0 um and 3.0-200 um) had the most mapped reads corresponding to a higher GV genome abundance in this size fraction. The smallest size ranges (0.1-0.8 um and 0.2-3 um) had the least mapped reads, showing most of these GVs to be cell-associated with larger hosts. Regression analysis showed a significant positive relationship between sequencing depth and genome recovery (p =0.0014) with no genomes recovered below 25 million paired-end reads. The sample with the most recovered genomes had a sequencing depth of ∼87 million paired-end reads.

**Figure 7.**
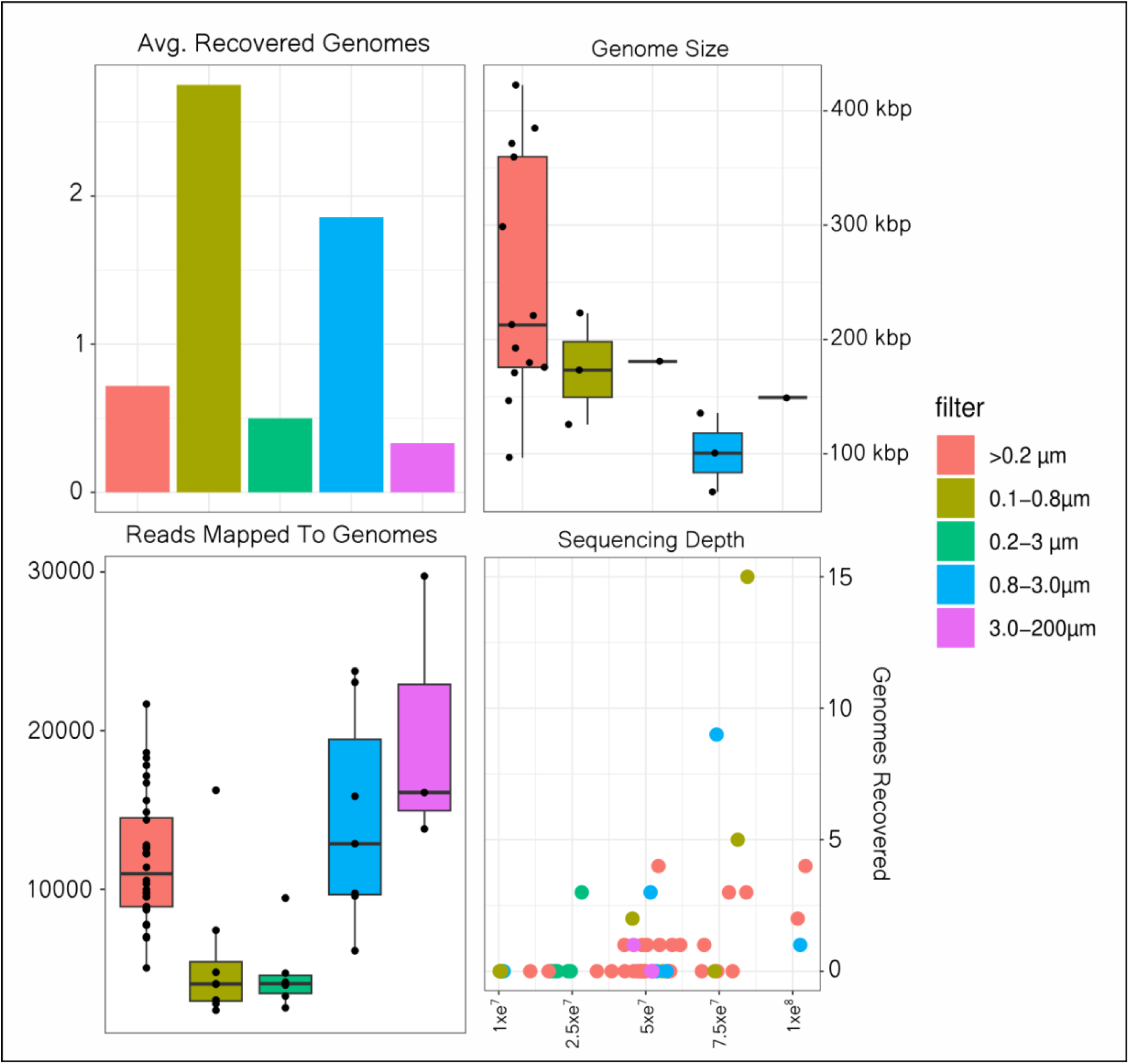
A case study on factors affecting GVMAG recovery. Samples from the Baltic Sea (n=56) were selected to gain insight into how filter fraction and sequencing depth play a role in GVMAG recovery. Reads from these samples were also mapped to Baltic Sea GVMAGs to see relative abundance in each filter fraction. Reads were normalized for library size.

### Environmental factors associated with Baltic Sea GV community composition

A Mantel test and CCA analysis were used to determine environmental factors driving GV community composition in the Baltic Sea. When all GVs were considered together, bacterial production, depth, and salinity were predicted to be significantly correlated with GV abundance (p <.05) (Figure 8a). Genomes were also separated by order level classification to see specific phylogenetic differences in factors affecting community composition. Genomes belonging to the Algavirales and Imitervirales orders were only significantly correlated with depth. Asfuvirales were positively correlated with dissolved organic nitrogen and dissolved organic carbon. Pandoravirales were only correlated with bacterial production, and Pimascovirales were correlated with bacterial production, dissolved oxygen, and salinity. The CCA analysis confirmed salinity, bacterial production, and depth as the driving factors of the combined community (Figure 8b).

**Figure 8.**
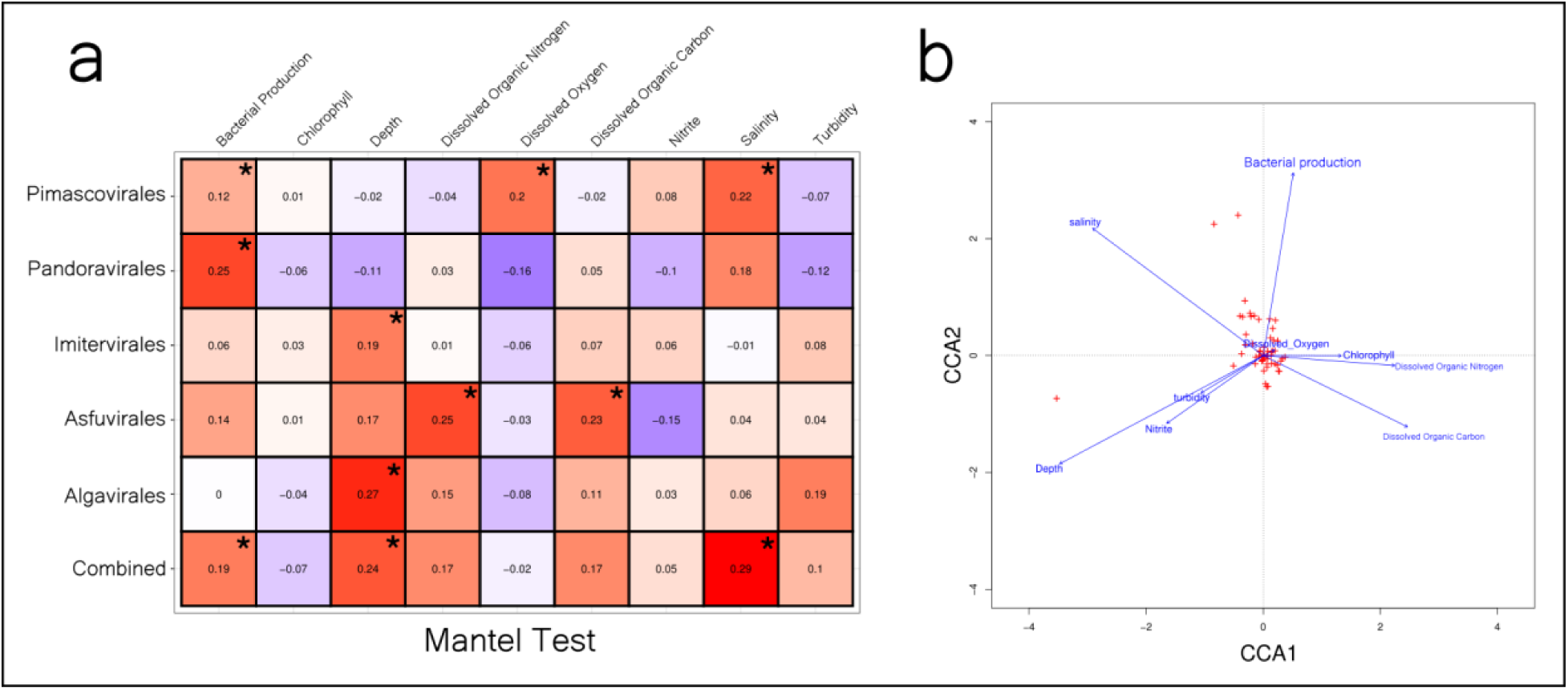
The correlation of environmental parameters with GVMAG abundance. Both a Mantel test and (b) CCA analysis were performed to see correlations with GVMAG abundance in the Baltic Sea. The Mantel test was conducted on all combined GVMAGs as well as each order that had more than 2 recovered GVMAGs, and the correlation coefficient is displayed (* = p<.05).

## Discussion

In an era of metagenomic research, giant virus diversity and functional potential have greatly been spurred on by a number of recent large-scale GV genome recovery approaches from publicly available metagenomic datasets [41–43]. These sweeping studies greatly increased the number of genomes available for GVs, but much work is still to be done to resolve their immense diversity in the marine ecosystem. Our 230 new genomes reported in this study provide additional resources for accessing GV diversity and functional potential. In addition to this, our easy-to-use pipeline, PIGv, should open up the field for further discovery of GVs in diverse datasets.

Analysis of our recovered genomes confirmed what was known broadly about differences between GV orders such as the higher GC percentage of Pandoravirales genomes (>60%) [3] compared to other orders [66]. Other metrics such as Nitrogen and Sulfur content of GV genomes have never been reported. The proteomic nitrogen stores have been hypothesized to give insight into the connection between ecology and genomics as lower nitrogen content can be a proteomic signature of environmental nitrogen scarcity [67]. In the marine environment, it is estimated that minimization of proteomic nitrogen can reduce the cellular nitrogen budget by up to 10% for marine microbes [52]. While viruses don’t directly have a nitrogen budget, during infection they have the potential to shift the host’s nitrogen budget through its own replication. Changing the host’s genomic and proteomic pool could be yet another way that these viruses influence host metabolism and survival in the ocean as GVs with higher N per sidechain could have a greater toll on the host nitrogen budget [17].

The findings of phylogeny-associated metabolic potential were largely consistent with previous studies looking at GV metabolic potential. Moniruzamman et al. [42], Ha et al. [5], and Farzad et al. [66] report the largest number of metabolic genes associated with the Imitervirales order, with the Algavirales order having much less DNA processing and nutrient transport genes. We also observed the scarcity of rhodopsins inside the Algavirales order consistent with these studies. Many of the metabolic genes are functional during infection. For example,MutS and HAT proteins were shown to be packed inside the capsid [68,69] and the presence of these genes more frequently in the Imitervirales order could imply a phylogenetic pattern in DNA packaging strategy within *Nucleocytoviricota* [70].

In addition to more metabolic genes, we also found the functional landscape between Imitervirales and Algavirales genomes to be quite different with Imitervirales genomes encoding a large array of genes not found in Algavirales. The repeated finding of more auxiliary metabolic genes as well as the broad functional capacity within the genomes of Imitervirales could be indicative of a different “life strategy” as Imitervirales viruses most likely infect a broad range of hosts including slow-growing hosts where metabolic augmentation may be necessary for productive infection [71].

In addition to uncovering known GV-encoded functional patterns, we found many proteins previously unknown to GVs to be found in our new high-quality and partial GVMAGs. This finding shows that the protein universe for GVs remains largely open as large amounts of novel proteins and functional capacity remain to be discovered, expanding our understanding of what functions virus genomes can encode and perform. A large number of these novel functions coming from polar or Baltic Sea GVMAGs could be further evidence of environmentally specific infection strategies or functional repertoires as is seen in other nutrient and light regimes [72,73]. Previous analysis of polar giant viruses also found high levels of adaptation and unique gene content compared to temperate and tropical counterparts [74], showing these colder environments could be reservoirs of novel encoded virus functions.

A well-explored and interesting area of GV functional potential is the ability of GVs to influence host photosynthetic potential through the introduction of rhodopsins and light-harvesting proteins [75–77]. Here, we show that not only are these genes present in our new genomes, but additional genes, involved in all stages of photosynthetic light harvesting, are present as well. It is well known that some GV infections depend on light and functioning photosynthetic machinery during the early stages of infection [78,79], so GVs may be using auxiliary genes to keep host machinery operational during infection as the host tries to shut off its transcriptional machinery[80,81]. Here we report many new proteins involved in the photosynthetic pathway, adding evidence to the claim that GVs can modulate this pathway in their host.

Within the context of the Baltic Sea ecosystem, recovered GVMAGs demonstrated differential abundance correlating with various environmental parameters. Previous work on prokaryotic viruses has shown the high explanatory power these parameters can have on viral abundance across coastal waters [82], and our data confirm this conclusion as factors such as salinity and depth showed significant correlation with GV abundance. Despite these factors being significant for the total GV population, factors affecting GV community composition differed by phylogeny, possibly reflecting the different hosts of viruses within these phylogenetic orders [9].

### Best practices for giant virus genome recovery

From our case study on the Baltic Sea samples, we saw some general trends toward boosting the recovery of GVs from metagenomic datasets. From the data available to us, it seemed the best size range for GV recovery was in the 0.1-0.8 uM fraction. The use of an upper constraint in filtering is a common trend in GV recovery in the marine environment as most GV genomes from the Tara Oceans data come from a size fraction with an upper constraint [43]. This upper constraint allows for the exclusion of many larger protists and organisms that would occupy a large portion of the sequencing space, thus maximizing GV recovery. Sequencing depth is also a question to consider when looking to recover GVMAGs. Using the Baltic data, it seemed that the minimum sequencing depth of 25 million paired-end reads was necessary for recovery, with incremental gains for higher sequencing depths. This is not a hard cutoff as genomes were recovered from Jangmok Bay sequencing data that had an average sample depth of 10 million paired-end reads. There did seem to be an upper limit at around 100 million paired-end reads where additional depth did not yield additional recovery. Overall, the usage of an upper constraint in the 0.8-3um range as well as a sequencing depth greater than 25 million paired-end reads is advised for recovery of GV genomes from metagenomic datasets.

## Conclusion

Overall, our work provides new insights into the diversity and functional potential of GVs in the world’s oceans through the introduction of a new pipeline to recover high-quality genomes from metagenomic datasets. Our addition of 230 genomes and new functional potential expands the knowledge base of GVs and their role in the marine ecosystem. We hope that our new pipeline along with methodological recommendations will be useful in the recovery of GVs from further metagenomic datasets across all aquatic ecosystems.

## Data Availability

All metagenomes used for this analysis are publicly available on NCBI using the project numbers in Table S1. The GVMAGs, Genome statistics, taxonomy, phylogenetic tree, PolB proteins, Anvi’o database, and protein annotations are available on figshare (https://figshare.com/projects/Expansion_of_the_genomic_and_functional_diversity_of_global_ocean_giant_viruses/214120).

## Acknowledgments

This study leveraged a large number of publicly available metagenomic datasets, and we thank the researchers who have made their data available. We also thank the researchers who made their bioinformatic tools and pipelines available for public use. We also gratefully acknowledge the computational resources provided by the Frost Institute for Data Science and Computing (IDSC), University of Miami.

## Author contributions

BM and MM jointly developed the research idea. BM performed the data analysis, bioinformatic pipeline development and wrote the manuscript. MM supervised the research, contributed writing and editing of the manuscript.

