## Supplemental Figures and Tables for "Expansion of the genomic and functional diversity of global ocean giant viruses"

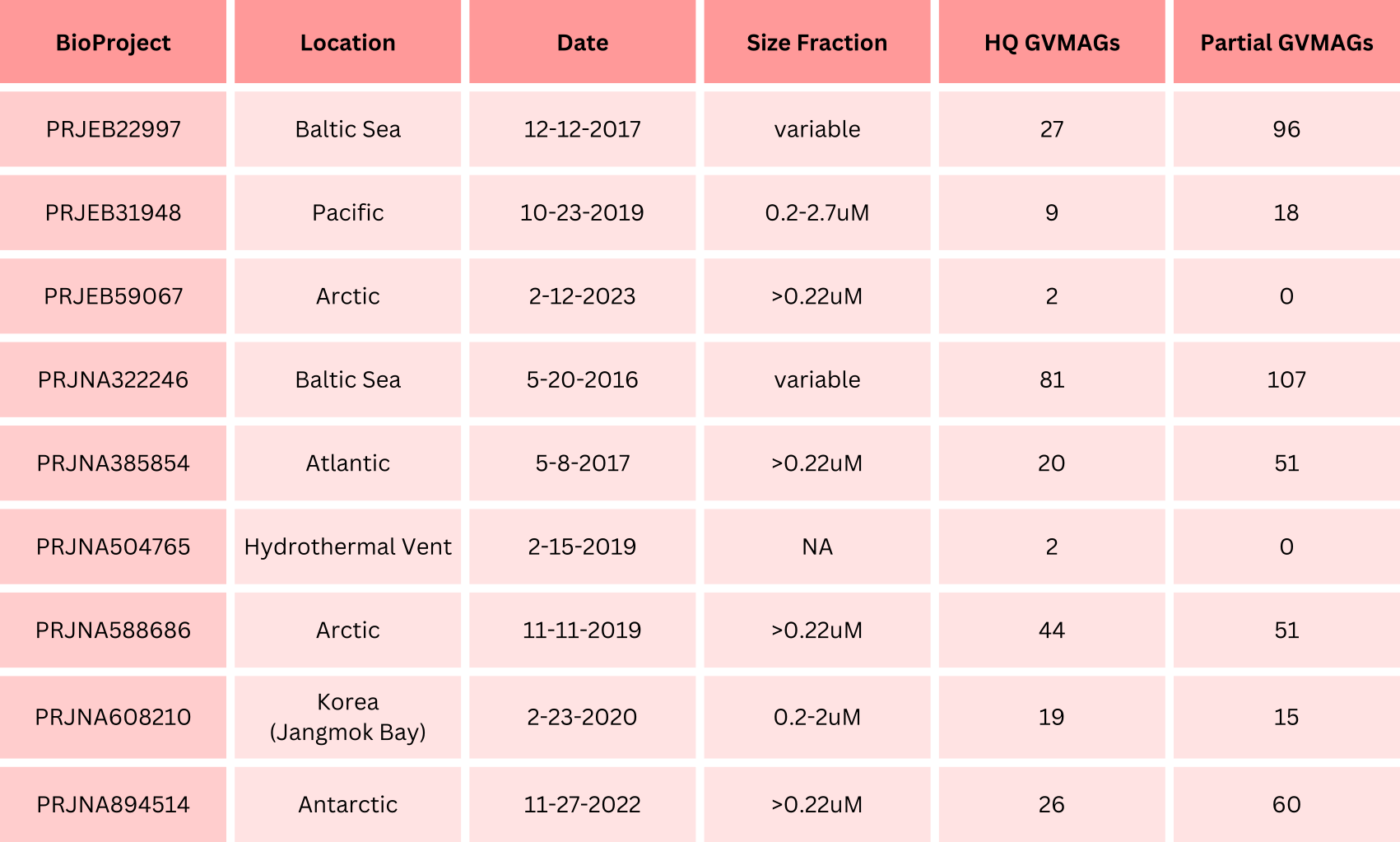

**Table S1. Studies used for this project**. Publicly available metagenomes were downloaded from NCBI from the listed BioProjects. The location of the study, date of data publication, size fraction used for filtering, as well as recovered genomes are presented in this table.

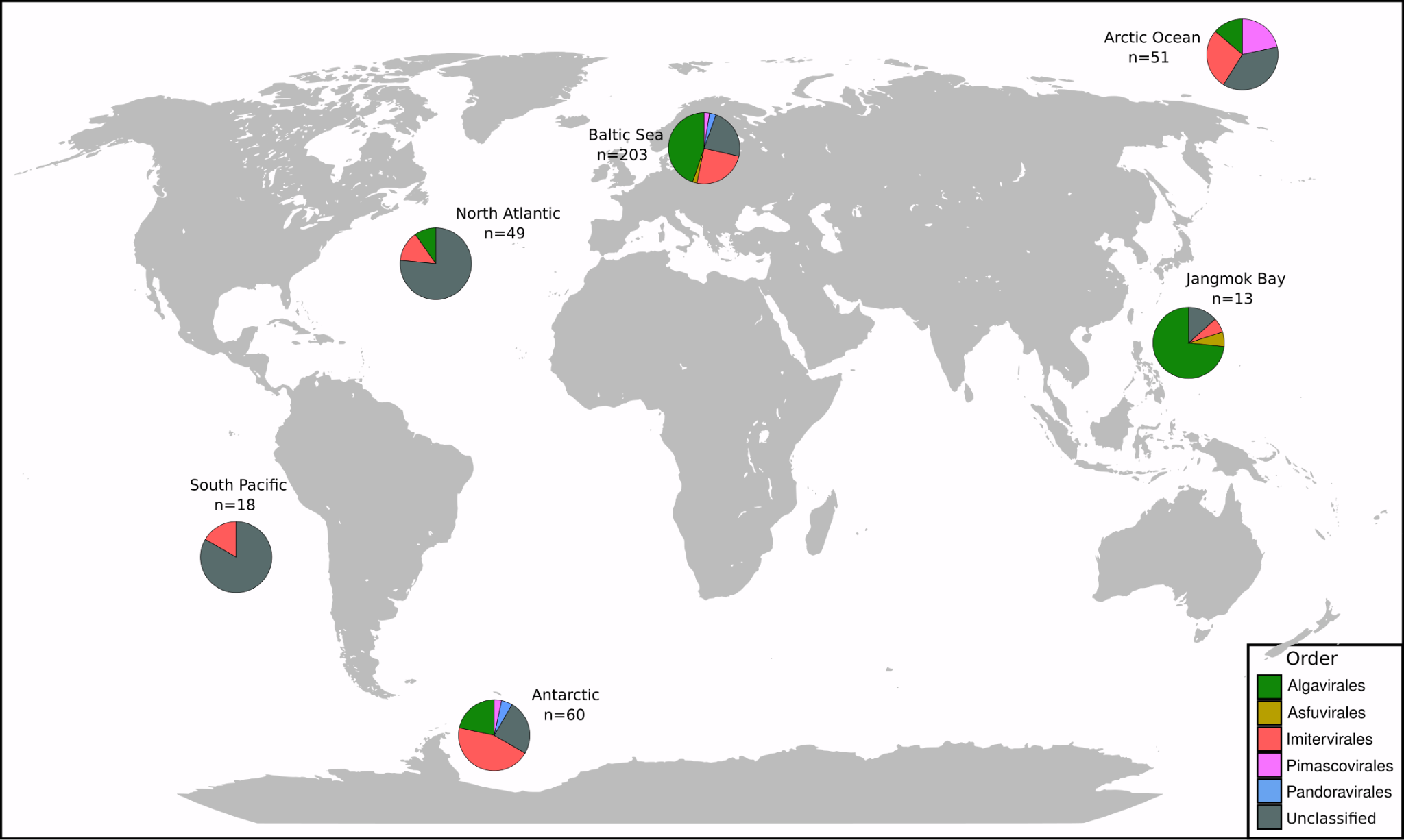

**Figure S1. Distribution of Partial Genomes**. A map showing the distribution of partial GVMAGs in the datasets. GVMAG classifications were made using TIGTOG.
